# A head-to-head comparison of two DREADD agonists for suppressing operant behavior in rats via VTA dopamine neuron inhibition

**DOI:** 10.1101/2023.03.27.534429

**Authors:** Kate A Lawson, Christina M Ruiz, Stephen V Mahler

**Affiliations:** Department of Neurobiology and Behavior, University of California Irvine, Irvine, CA USA

**Keywords:** chemogenetics, dopamine, ventral tegmental area, clozapine-n-oxide, JHU37160

## Abstract

**Rationale:** Designer receptors exclusively activated by designer drugs (DREADDs) are a tool for “remote control” of defined neuronal populations during behavior. These receptors are inert unless bound by an experimenter-administered designer drug, most commonly clozapine-n-oxide (CNO). However, questions have emerged about the suitability of CNO as a systemically administered DREADD agonist.

**Objectives:** Second-generation agonists such as JHU37160 (J60) have been developed, which may have more favorable properties than CNO. Here we sought to directly compare effects of CNO (0, 1, 5, & 10 mg/kg, i.p.) and J60 (0, 0.03, 0.3, & 3 mg/kg, i.p.) on operant food pursuit.

**Methods:** Male and female TH:Cre+ rats and their wildtype (WT) littermates received cre-dependent hM4Di-mCherry vector injections into ventral tegmental area (VTA), causing inhibitory DREADD expression in VTA dopamine neurons in TH:Cre+ rats. Rats were trained to stably lever press for palatable food on a fixed ratio 10 schedule, and doses of both agonists were tested on separate days in a counterbalanced order.

**Results:** All three CNO doses reduced operant food seeking in rats with DREADDs, and no CNO dose had behavioral effects in WT controls. The highest tested J60 dose significantly reduced responding in DREADD rats, but this dose also *increased* responding in WTs, indicating non-specific effects. The magnitude of CNO and J60 effects in TH:Cre+ rats were correlated and were present in both sexes.

**Conclusions:** Findings demonstrate the usefulness of directly comparing DREADD agonists when optimizing behavioral chemogenetics, and highlight the importance of proper controls, regardless of the DREADD agonist employed.

## Introduction

Designer receptors exclusively activated by designer drugs (DREADDs) are a useful method for attaining “remote control” of neuronal populations (Armbruster et al., 2007; Rogan and Roth, 2011). These mutated receptors are derived from human muscarinic acetylcholine receptors but are not affected by endogenous neurotransmitters, rendering them normally inert when expressed in targeted neural populations. Yet when a drug capable of agonist binding to DREADDs is administered experimentally, the receptors engage endogenous G protein-coupled signaling pathways, resulting in net excitation or inhibition of neuronal firing, and/or neurotransmitter release (Alexander et al., 2009; Atasoy and Sternson, 2018; Brodnik et al., 2020; Buchta et al., 2017; Mahler et al., 2019, 2014; Martinez et al., 2023; Song et al., 2022; Stachniak et al., 2014). Especially when coupled with genetic strategies such as cre-driver rodent lines, DREADDs are a powerful tool for manipulating G protein-coupled receptors in phenotypically-defined neural populations, elucidating the consequences of neural manipulations on behaviors and other outcomes (Burnett and Krashes, 2016; Ferguson et al., 2013; Fortress et al., 2015; Mazzone et al., 2018; O’Neal et al., 2020; Rinker et al., 2017; Rorabaugh et al., 2017; Zhu and Roth, 2015).

One of the most useful features of DREADDs is their “lock and key” nature—the premise that DREADDs (the “lock”) are inert in the absence of an exogenous ligand (the “key”), and that their ligand, when administered, acts “Exclusively” at DREADD receptors. Yet recent evidence suggests that CNO does not efficiently penetrate the blood brain barrier, and it binds relatively weakly at DREADDs. Instead, it is likely that metabolic conversion of CNO to clozapine, which enters the brain efficiently and binds DREADDs potently, is directly responsible for behavioral effects of systemically administered CNO. In other words, CNO essentially acts as a pro-drug for clozapine, the direct DREADD agonist. Since at high enough concentrations clozapine can also bind endogenous receptors, it is therefore also possible for CNO to have off-target effects (Gomez et al., 2017; Ilg et al., 2018; MacLaren et al., 2016; Manvich et al., 2018; Porter et al., 2017). CNO is frequently found to have no measurable behavioral effects in the absence of DREADDs (Mahler and Aston-Jones, 2018; Smith et al., 2016; Urban and Roth, 2015; Whissell et al., 2016), but other experiments have found non-specific effects (Bonaventura et al., 2019; Gomez et al., 2017; MacLaren et al., 2016; Manvich et al., 2018; Martinez et al., 2019; Porter et al., 2017; Raper et al., 2017). The reasons for these varying results are unknown, but may involve species differences, the behavioral task tested, the presence of other experimentally-administered drugs, and CNO dose, route of administration, and dosing frequency (Campbell and Marchant, 2018; MacLaren et al., 2016). It is thus essential to compare CNO effects in subjects with and without DREADDs, to determine specificity of observed behavioral changes to manipulation of the targeted neural population.

Regardless, there is a clear need for a next generation of DREADD agonist that binds potently and selectively to DREADDs at doses in which it has no off-target behavioral effects. Several groups have proposed alternatives to CNO for behavioral chemogenetic experiments. One strategy is to administer low doses of clozapine, which binds at lower doses to DREADDs than at endogenous receptors (Desloovere et al., 2021; Jendryka et al., 2019). Other candidate drugs with varying advantages and disadvantages include deschloroclozapine (DCZ), C21, olanzapine, and JHU37160 (J60) (Bonaventura et al., 2019; Desloovere et al., 2021; Ferrari et al., 2022; Fleury Curado et al., 2021; Goutaudier et al., 2020; Kljakic et al., 2022; Nagai et al., 2020; Nentwig et al., 2022; Thompson et al., 2018).

One of these compounds, J60, was developed by an NIH team who showed it has acceptable behavioral and other effects in mice (Bonaventura et al., 2019), a finding that has been replicated by several other groups (Desloovere et al., 2021; Flerlage et al., 2022; Fleury Curado et al., 2021; Giannotti et al., 2021; Heinsbroek et al., 2021; Huang et al., 2021; Lewis et al., 2020; Li and Hollis, 2021; Salimi-Nezhad et al., 2023; Zhang et al., 2020). J60 efficiently crosses the blood-brain-barrier, it binds directly to central DREADD receptors after i.p. administration, in mice it is behaviorally-effective at both excitatory (hM3Dq) and inhibitory (hM4Di) DREADDs, and it also shows promise in primates (Bonaventura et al., 2019). We also showed in the same report that it is effective at stimulating locomotor activity at hM3Dq excitatory DREADDs in VTA dopamine neurons of TH:Cre+ rats, even at very low doses that had no clear off-target actions (Bonaventura et al., 2019). One group showed 0.1 mg/kg J60 in rats has specific behavioral effects at hM4Di DREADDs, but behavioral effects of other doses with inhibitory DREADDs have not yet been reported in rats (Giannotti et al., 2021; Heinsbroek et al., 2021).

In general, there is a notable lack of head-to-head behavioral comparisons of DREADD agonists, delaying the field from advancing toward consensus on the best compound for use in behavioral neuroscience experiments. Toward this goal, we conducted a preliminary experiment comparing CNO to J60 in tyrosine hydroxylase (TH):Cre rats with inhibitory hM4Di DREADDs in ventral tegmental area (VTA) dopamine neurons. We tested multiple doses of each compound in the same animals, examining effects on performance of an instrumental task—pressing a lever on a fixed-ratio (FR)10 schedule for palatable food pellets. Our results indicate that high doses of both CNO and J60 similarly suppressed instrumental responding in rats with inhibitory DREADDs, though the highest dose of J60 tested had the notable paradoxical effect of enhancing lever pressing in control rats without DREADDs. These data support the idea that either of these compounds can be used in such behavioral neuroscience experiments, though careful dosing optimization, and direct comparison of effects in animals with and without DREADDs is required. We also hope this report will inspire further direct comparisons of DREADD agonists to one another across different classes of behaviors and ranges of doses.

## Methods

All procedures were approved by the Institutional Animal Care and Use Committee at UC Irvine, and are in accordance with the NIH Guide for the Care and Use of Animals.

### Subjects

Long Evans transgenic TH:Cre+ (N=13) and wildtype TH:Cre- (WT; N=9) rats (N=9 males, N=13 females) were bred in-house, and housed as adults in pairs in ventilated tub cages with corncob bedding and *ad libitum* chow and water. Rats were at least 75 days old at the start of experiments. Rats were housed in reverse 12:12 hr lighting, and behavior experiments took place during the dark cycle.

### Drugs

CNO was provided by the NIDA Drug Supply Program, stored in desiccated, opaque powder aliquots at 4 °C, and prepared daily, mixed in 5% dimethyl sulfoxide (DMSO) in saline solution. J60 was provided by the NIMH Drug Supply Program, stored in desiccated, opaque powder aliquots at 4 °C, and prepared weekly, mixed in 5% DMSO saline solution and also stored at 4°C.

### Viral Vector and Surgery

A validated (Mahler et al., 2019, 2014) AAV2-hSyn-DIO-hM4D(Gi)- mCherry vector (titer ≥ 5×10¹² vg/mL) was attained from AddGene (catalog number 44362-AAV2). Rats were anesthetized with 2.5% isoflurane, with meloxicam analgesic (1.0 mg/kg), then stereotaxically injected via glass pipette and Picospritzer with 0.75 µl of the vector bilaterally, aimed at the VTA (coordinates relative to bregma (mm): -5.5 AP, +/- 0.8 ML, −8.1 DV). Pipettes were left in place for 5 min to reduce spread prior to removal. Rats were allowed at least 10 days to recover following surgery before beginning training. At least 28 days elapsed between virus injection and the first administration of CNO or J60, allowing sufficient time for robust, persistent DREADD/reporter expression in VTA dopamine neurons (Brodnik et al., 2020; Mahler et al., 2019, 2014).

### Behavioral Training and Testing

After recovering from surgery, animals were trained to lever press for highly palatable, banana-flavored, sucrose, fat, and protein-containing pellets (Bio-Serv, catalog #F0059) in a Med Associates rat operant conditioning box, enclosed in a sound-proof chamber. In daily 1 hr sessions, rats began on a fixed ratio (FR) 1 schedule and moved up to FR3, FR5 then FR10 when their responding was consistent for 3 consecutive days. No genotype or sex difference was seen in the number of days to progress to FR10, or to stabilize on FR10 responding (ps>0.05). After at least 8 days at FR10, and when stability criterion was achieved (less than 33% change in responding for 2 consecutive days), testing with DREADD agonists commenced. Animals were tested in a counterbalanced, pseudo-random order, with tests of all CNO doses (vehicle, 1, 5, 10 mg/kg) conducted prior to beginning a counterbalanced series of J60 tests (vehicle, 0.03, 0.3, 3 mg/kg). Stable responding was re-established, and >48 h elapsed between tests. The two vehicle (VEH; 5% DMSO in saline solution) test days were not statistically different from one another (p>0.5), so they were averaged for further analyses and in figures. Experimental timeline is shown in **Fig. 1**.

**Fig. 1.**
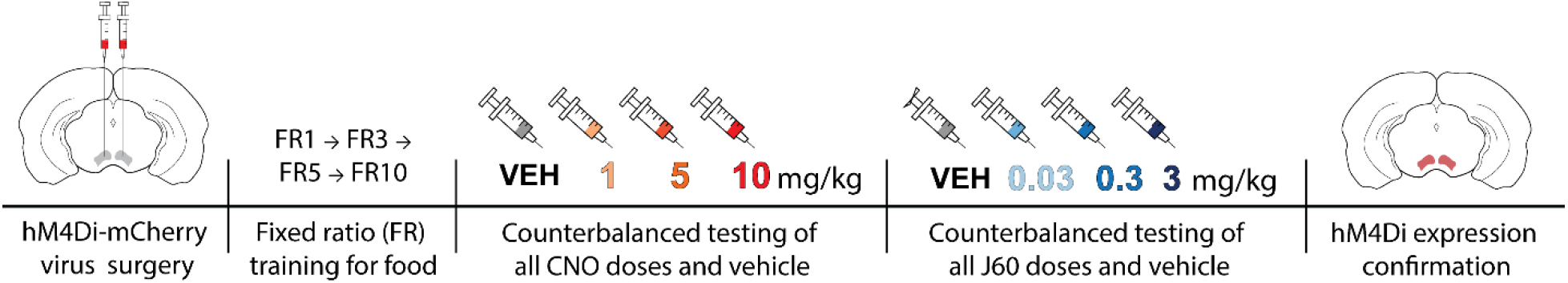
Schematic of experimental timeline. Following hM4Di DREADD injection into the VTA, rats underwent fixed ratio (FR) training for palatable food and were stably responding at FR10 before testing with DREADD agonists began. All CNO doses and a VEH test were counterbalanced, followed by counterbalanced J60 doses and another VEH test. Animals were sacrificed after all tests were completed to confirm DREADD expression

### Confirmation of DREADD Expression

After the final test was completed, animals were transcardially perfused with ice cold 0.9% saline followed by 4% paraformaldehyde. Extracted brains were cryoprotected in 20% sucrose, sectioned at 40 µm in a cryostat, and blocked in 3% normal donkey serum PBST. Tissue was incubated for 16 hr in rabbit anti-DsRed (Clontech; 1:5000) and mouse anti-TH antibodies (Immunostar; 1:1000) in PBST-azide with 2% normal donkey serum. After washing, slices were incubated in the dark for 4 hr in AlexaFluor-donkey anti-rabbit 594 and donkey anti-mouse 488 (Thermo Fisher Scientific), washed, then incubated with 4′,6-diamidino-2-phenylindole (DAPI; 1:1000) in PB for 5 mins, washed, mounted, and coverslipped with Fluoromount (Thermo Fisher Scientific). mCherry, TH, and DAPI expression was imaged at 10x, and the zone of expression in each hemisphere of each rat was verified relative to VTA borders using a rat brain atlas (Paxinos and Watson, 2006) (**Fig. 2**). Colocalization of TH and mCherry was also visualized at 63x magnification, and showed specific expression of mCherry in VTA dopamine neurons, as previously reported (Mahler et al., 2019, 2014).

**Fig. 2.**
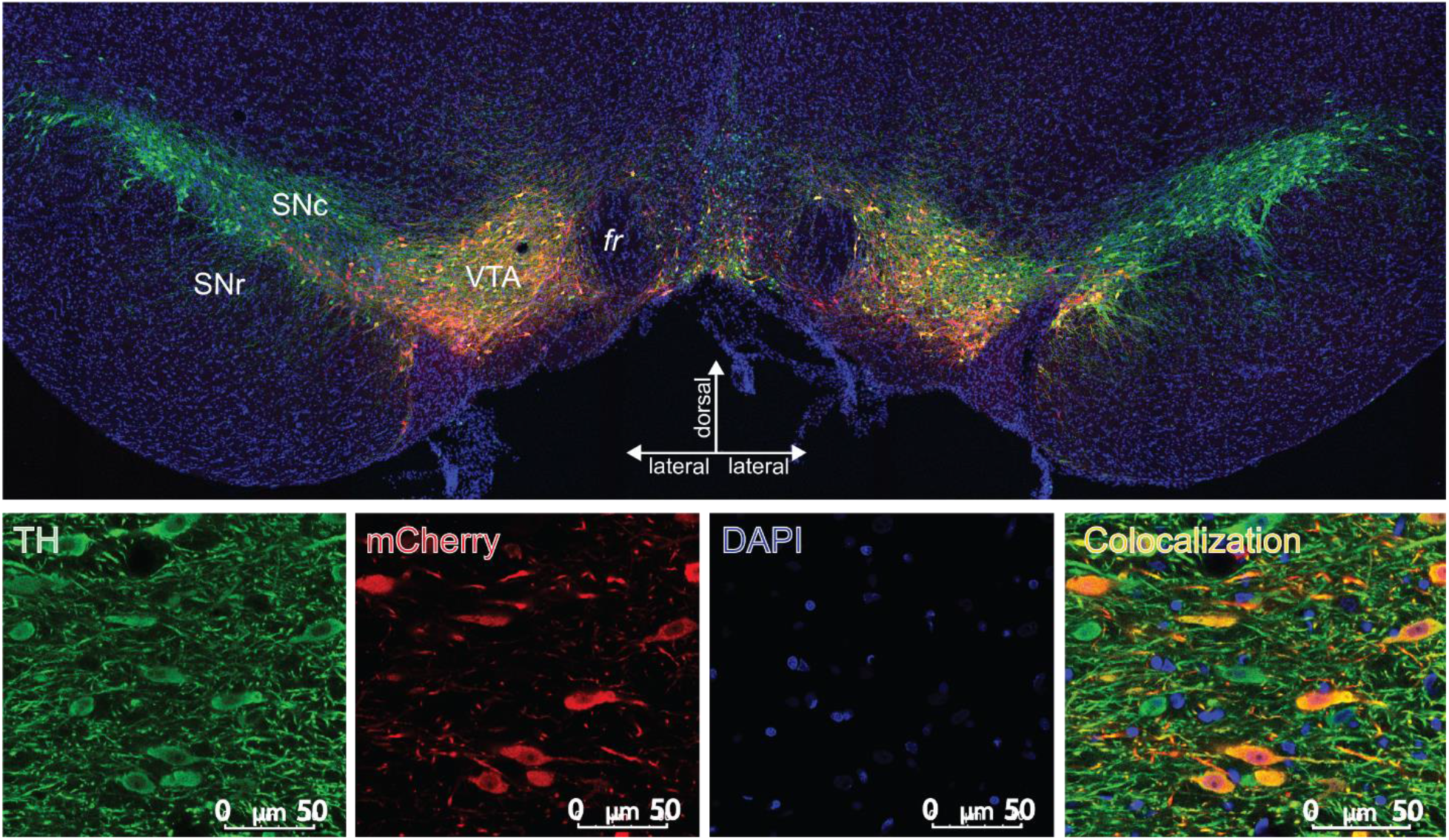
Bilateral VTA Dopamine Neuron DREADD Expression: (Top) Typical expression of AAV2-hSyn-DIO-hM4D(Gi)-mCherry vector in a TH:Cre+ rat is depicted in a coronal view. mCherry (the cre-dependent DREADD reporter; red stain) is expressed nearly exclusively in TH+ neurons (green stain) of VTA; DAPI counterstain (blue). (Bottom) Each stain is shown separately at higher magnification (Scale bar=50 µm)

### Statistical Analyses

Analysis of drug effects was conducted on change from baseline data, since rats’ baselines could drift over the course of training. We averaged active lever presses, inactive lever presses, and rewards earned on the 2 days prior to the test day and subtracted that average from test day values. We performed two-way ANOVAs on each drug, with dose (within subjects, CNO: vehicle, 1, 5, and 10 mg/kg; J60: vehicle, 0.03, 0.3, 3 mg/kg) and genotype (between subjects, TH:Cre+ and WT) factors. Separate ANOVAs were used to analyze active and inactive lever presses. Significant ANOVAs were followed up with one-way ANOVAs comparing doses in TH:Cre+ and WT rats, with Tukey HSD posthoc tests. When sex was added to ANOVAs, there was no main effect of sex, nor any interactions of sex with other variables, so sexes were combined for subsequent analyses. Pearson’s correlation was used to determine whether behaviorally inhibitory effects of CNO and J60 were correlated (change from baseline vehicle day responding subtracted from change from baseline CNO/J60 dose responding; [test day – 2 d prior baseline] – [vehicle day – 2 d prior baseline]). In all cases, two-tailed tests with significance thresholds of p>0.05 were used. Statistical analyses were conducted in R.

## Results

### Viral Expression

TH:Cre+ rats exhibited hM4Di-mCherry expression that was localized within VTA borders, and expression was observed to be highly selective to dopamine neurons, as previously described (**Fig. 2**) (Brodnik et al., 2020; Mahler et al., 2019, 2014). Two TH:Cre+ animals were excluded from behavioral analyses because they lacked mCherry expression in the VTA in one hemisphere, and one animal was excluded due to a broken operant lever during testing, for a total of 11 TH:Cre+ and 8 WT animals included in analyses.

### Effects of CNO on FR10 Responding

CNO inhibited active lever pressing relative to baseline selectively in TH:Cre+ rats (CNO dose X genotype interaction on change from pre-test baseline responding: F_3,51_=3.88, p=0.0142; **Fig.3A**). In TH:Cre+ animals, there was a main effect of dose (F_3,30_=9.64, p=0.000129), which was driven by suppression of responding, relative to vehicle, at each tested dose (1 mg/kg: p=0.0446; 5 mg/kg: p=0.00495; 10 mg/kg: p=0.00481). There was no main effect of CNO doses in WT animals (F_3,21_=0.147, p=0.930; **Fig.3C**), so CNO only inhibited palatable food pursuit in animals with hM4Di DREADDs.

**Fig. 3.**
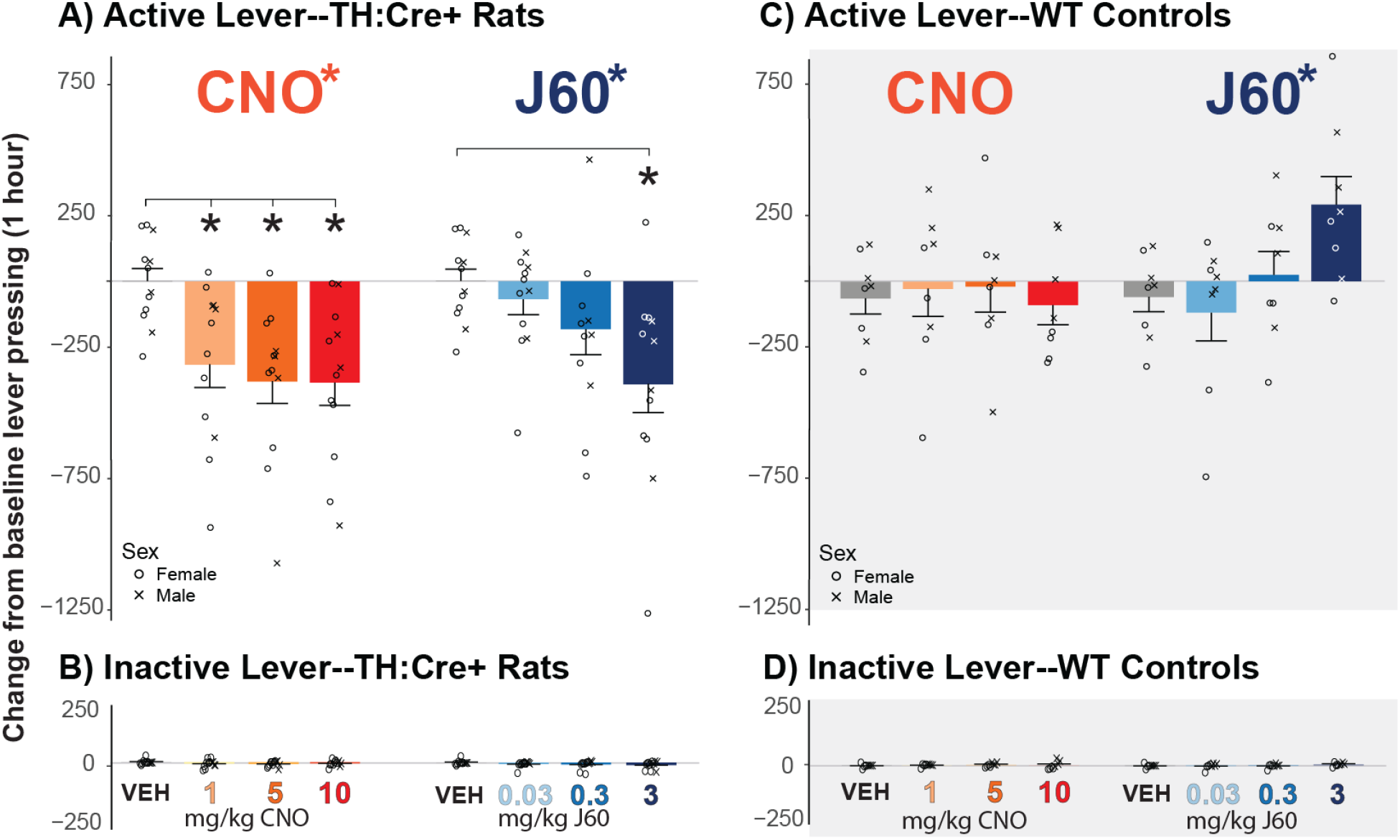
CNO and J60 Effects on Operant FR10 Responding. A) Change in active lever presses from 2 day prior average baseline with administration of CNO and J60 in TH:Cre+ animals is shown. There was a main effect of both CNO and J60 on active lever presses in TH:Cre+ animals, with posthoc test results comparing each dose to vehicle indicated with *; p<0.05). B) Change in inactive lever presses from baseline in TH:Cre+ animals. C&D) Change in (C) active and (D) inactive lever pressing is shown for WT animals. There was a main effect of J60 on active lever pressing in WT animals. Crosses and circles depict data from individual male and female animals, respectively

CNO similarly suppressed the baseline-relative number of rewards earned in TH:Cre+, but not WT rats (dose X genotype interaction: F_3,51_= 4.50, p=0.00705). In TH:Cre+ rats, there was a main effect CNO dose (F_3,30_=9.95, p=0.000103), driven by a drop in rewards earned between VEH and each CNO dose (1 mg/kg: p=0.0446; 5 mg/kg: p=0.00495; 10 mg/kg: p=0.00480).There was no effect of CNO on rewards earned in WT animals (F_3,21_=0.233, p=0.872).

### Effects of J60 on FR10 Responding

J60 also had distinct, dose-dependent effects on change from baseline active lever pressing in TH:Cre+ and WT rats (genotype X dose interaction: F_3,51_=9.23, p=0.0000556). In TH:Cre+ rats, there was a main effect of dose (**Fig. 3A**; F_3,30_=5.67, p=0.00337), driven a drop in active lever pressing between VEH and J60 3 mg/kg (p= 0.00241). However there was also an effect of dose in WT animals (**Fig. 3C**; F_3,21_=4.20, p=0.0178), and though there were no significant changes from VEH at any dose in posthoc analyses, the high dose trended toward increasing responding (3 mg/kg: p=0.0551).

JHU had similar effects on number of rewards earned as it did on active lever responding (genotype X dose interaction (F_3,51_=9.40, p=0.0000477). In TH:Cre+ rats, there was a main effect of dose (F_3,30_=6.60, p=0.00147), driven by a drop in rewards earned between VEH and J60 3 mg/kg (p=0.00139). There was also an effect of dose in WT animals (F_3,21_=3.79, p=0.0257), driven by a significant increase in food pellets earned at the high dose (3 mg/kg: p=0.0492).

### Inactive Lever Responding

There were no significant effects of either CNO or J60 on inactive lever pressing relative to baseline in either genotype (**Fig. 3B, 3D**; dose x genotype interaction; CNO: F_3,51_=1.25, p=0.300; J60: F_3,51_=1.76, p=0.166). Accordingly, in TH:Cre+ rats there was no effect of CNO (F_3,30_=1.56, p=0.218) or J60 (F_3,30_=1.32, p=0.285). Likewise, in WT rats there was no effect of either CNO (F_3,21_=0.178, p=0.910) or J60 (F_3,21_=1.16, p=0.349).

### Comparison of CNO to J60-Inhibited Responding

To determine whether the ability of CNO and J60 to suppress responding were of similar magnitude in individual animals, we next transformed change from baseline data for each dose to compute a change from vehicle score. This score was calculated by computing the change from baseline responding (average of 2 days prior) on each CNO or J60 dose, and subtracting from this the change from baseline on vehicle day. No statistical difference between the DREADD agonist drugs was observed in a dose (Low; Mid; High) X drug (CNO; J60) repeated measures ANOVA in TH:Cre+ rats (main effect of dose: F_2,20_=4.17, p=0.0307; but no main effect of drug: F_1,10_=3.04, p=0.112; or dose x drug interaction: F_2,20_=2.51, p=0.107).

To further query whether behavioral effects of chemogenetic VTA dopamine neuron inhibition with CNO versus J60 were related in individual rats, we next examined correlations between VEH-relative pressing after moderate (CNO: 5 mg/kg; J60: 0.3 mg/kg) or high doses of each drug (CNO: 10 mg/kg; J60: 3 mg/kg), in TH:Cre+ or WT rats. For the moderate doses, pressing suppression by both drugs trended toward correlation in TH:Cre+ rats (**Fig. 4A**; r=0.543, p=0.0843) but not WT littermates (**Fig. 4B**; r=-0.425, p=0.294). For the high doses, CNO- and J60-suppression of pressing was highly correlated in TH:Cre+ rats (**Fig. 4C**; r=0.762, p=0.00634), but no such effects were seen in WT littermates (**Fig. 4D**; r=-0.00916, p=0.982).

**Fig. 4.**
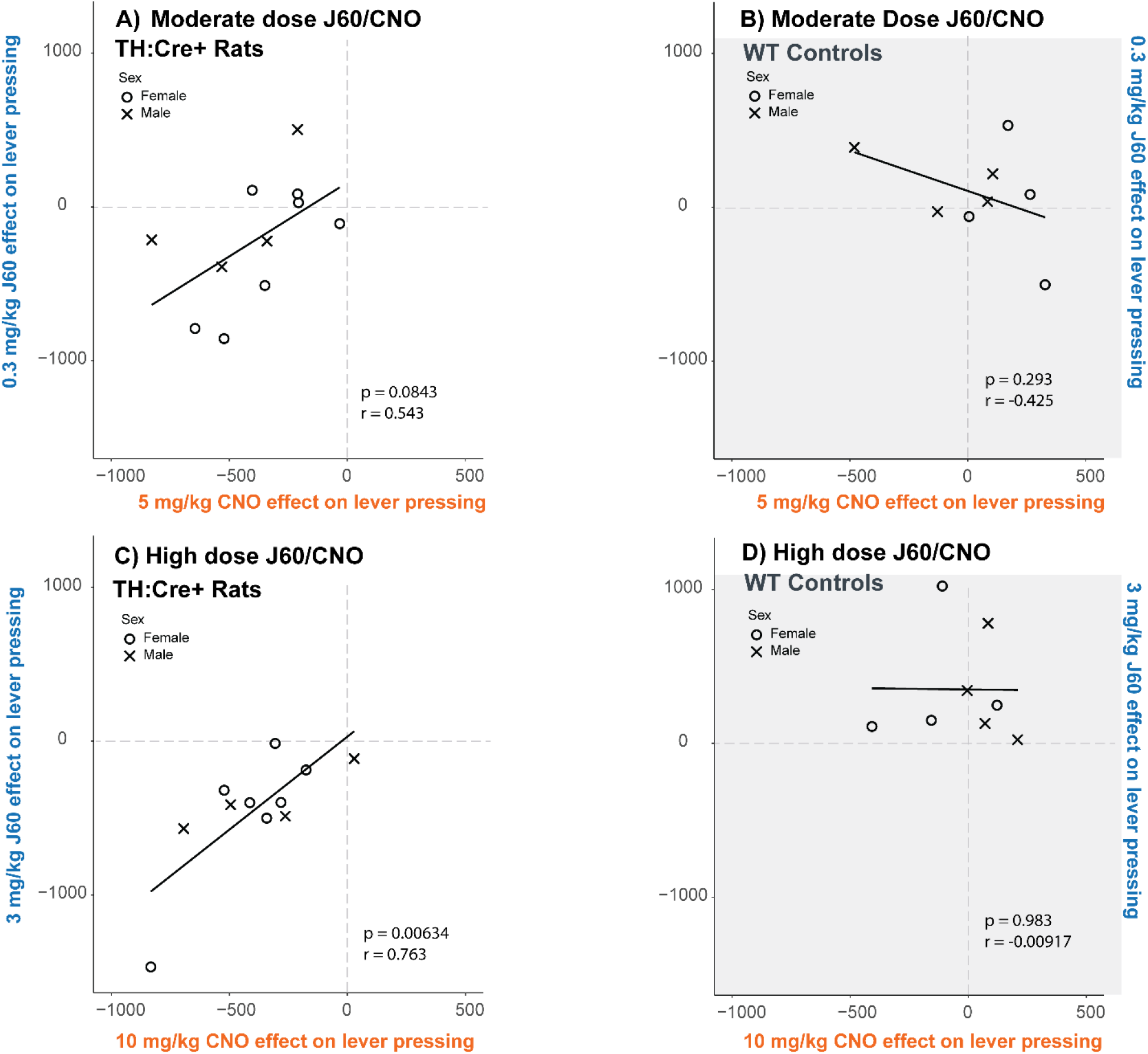
CNO and J60 Suppression of Pressing is Correlated in TH:Cre+, but not WT rats. A&B) Correlation between effects of moderate CNO and J60 doses on suppressing active lever pressing in A) TH:Cre+, and B) WT animals is shown. C&D) Correlation between effect of high dose CNO and J60 is shown in C) TH:Cre+ and D) WT animals. Data is shown as VEH- and baseline-relative. Crosses and circles depict data from individual male and female rats, respectively

## Discussion

DREADDs are a common approach for manipulating neural populations and circuits of behaving animals in neuroscience experiments. However, there remains controversy over which is the best agonist drug for engaging DREADDs. Therefore, we elected to test two prominent DREADD agonists (clozapine-n-oxide; CNO, and JHU37160; J60) head-to-head, using behaving TH:Cre+ rats expressing hM4Di inhibitory DREADDs in VTA dopamine neurons, or WT littermates without DREADDs. Using an operant reward seeking task (FR10 lever pressing for palatable food), we found that both agonists inhibited reward seeking and rewards obtained in hM4Di DREADD-expressing animals, and CNO did so at doses that did not affect behavior in WT controls. We also found that J60 *enhanced* reward attainment in WT rats at the highest dose (3 mg/kg), despite strongly suppressing seeking in TH:Cre+ rats at the same dose. The magnitude of CNO- and J60-suppression of reward across rats was also correlated in rats with VTA dopamine neuron hM4Di DREADDs, but not in WT controls. Taken together, these results suggest that both CNO and J60 can activate inhibitory DREADDs in VTA dopamine neurons to suppress operant food seeking. An important implication of these studies is that whatever agonist drug is used in a DREADD experiment, it is essential to compare its effects in experimental animals to effects in control animals without DREADD expression.

CNO significantly reduced active lever pressing in hM4Di DREADD rats at all tested doses (1, 5, 10 mg/kg) without having any behavioral effect in non-DREADD WT rats. J60 significantly reduced lever pressing in TH:Cre+ rats at the highest tested dose (3 mg/kg), but this dose also showed signs of increasing responding in WT animals, suggesting nonspecific effects. Supporting the qualitatively similar efficacy of CNO and J60, we found that the highest tested doses of both drugs elicited statistically equivalent behavioral effects in the same animals, and that the magnitude of effects elicited by these drug doses in individual animals was correlated.

We picked these specific doses of CNO based on precedent within our own lab and in the field more broadly. We’ve seen specific behavioral effects of CNO in rats at doses of 1, 5, 10, and even up to 20 mg/kg (Farrell et al., 2021, 2019; Lawson et al., 2021; Mahler et al., 2014). J60 has previously been tested at doses from 0.01 to 0.3 mg/kg with specific behavioral effects (Bonaventura et al., 2019; Desloovere et al., 2021; Giannotti et al., 2021; Heinsbroek et al., 2021). The high dose of J60 used here, 3 mg/kg, is likely the highest ever tested. At this dose, J60 effectively inhibits active lever pressing in animals with DREADDs, but also showed nonspecific effects of increasing reward obtainment in WT rats. It is not presently clear which receptor this high dose of J60, or its potential metabolites, might act at to produce these non-specific response-facilitating effects.

In contrast to J60, we did not observe any off-target effects of CNO at any tested dose. Some prior studies have found non-selective effects of CNO in DREADD-free rats or mice (Bonaventura et al., 2019; Desloovere et al., 2021; Gomez et al., 2017; Jendryka et al., 2019; MacLaren et al., 2016; Manvich et al., 2018; Porter et al., 2017; Raper et al., 2017), though we and many others have failed to find CNO-only effects on behavior in operant responding for food and drugs in our prior work (Farrell et al., 2021, 2019; Mahler et al., 2014). It is possible that the non-specific behavioral effects of CNO are dependent upon the behavior being tested. It is also possible that CNO effects are exacerbated by the presence of other drugs, potentially due to competitive metabolism of drugs that could enhance overall exposure to the agonist or its metabolites, such as clozapine (Mahler et al., 2019). Such metabolic competition may vary between species, strains, and sexes, and can also depend on the animals’ health—it is clearly a topic that requires further, dedicated study. Regardless, we strongly recommend that all DREADD studies using CNO or other agonists employ proper control groups to account for the potentially task-specific effects of CNO (or any DREADD agonist) in the absence of DREADDs.

### Limitations and Future Directions

These studies, testing in the same rats the relative efficacy of two common DREADD agonist drugs in eliciting hM4Di-dependent behavioral effects, leave several important questions unanswered. Instead of having the drug administration order fully counterbalanced, J60 doses were always given after CNO doses. The effects of J60 on behavior could therefore be impacted by prior CNO administrations, or repeated engagement hM4Di receptors by CNO. That being said, we have previously found that repeated CNO administrations in TH:Cre rats with hM4Di DREADDs in dopamine neurons did not have lingering effects on operant reward seeking (Farrell et al., 2019; Mahler et al., 2019, 2014), yet it is still possible that CNO-induced plasticity could have had subtle effects here that impacted effects of J60.

We tested 3 doses of each agonist drug, but these are not necessarily the optimal doses for controlling behavior via selective actions at hM4Di DREADDs. For example, while we observed a non-specific effect of high dose J60, it is possible that a dose between 0.3 and 3 mg/kg would have had strong behavioral effects on this task that were highly specific to DREADDs. Indeed, mg/kg J60 i.p. in rats has been reported to hM4Di DREADD-specifically suppress neural activity and alter drug seeking behavior (Giannotti et al., 2021; Heinsbroek et al., 2021), and both 0.1 and 1 mg/kg doses have been reported to effectively alter behavior and suppress neuronal activity at hM4Di DREADDs in mice (Bonaventura et al., 2019; Li and Hollis, 2021; Zhang et al., 2020). A group using both CNO and J60 found no differences between behavioral changes elicited by 0.1 mg/kg J60 and 3 mg/kg CNO at hM4Di DREADDs in mice (Lewis et al., 2020), and J60 is also effective at 0.1 and 1 mg/kg doses in rats and mice at hM3Dq DREADDs (Huang et al., 2021; Salimi-Nezhad et al., 2023; Zhang et al., 2020). These doses of J60 may therefore have selective behavioral effects depending on the task and neural substrate targeted, and may not be apparent in FR10 responding for palatable food. Further dose characterization with this new compound remains to be thoroughly characterized in both rats and mice, and future work with J60 should test doses for selectivity and efficacy in the behavioral model of interest.

In addition, other DREADD agonists are also promising, and should be similarly tested empirically. For example, compound 21 (Ferrari et al., 2022; Jendryka et al., 2019; Kljakic et al., 2022; Thompson et al., 2018), perlapine (Chen et al., 2015; Kljakic et al., 2022; Thompson et al., 2018), DCZ (Nagai et al., 2020; Nentwig et al., 2022; Raper and Galvan, 2022; Upright and Baxter, 2020), and olanzapine (Goossens et al., 2021; Upright and Baxter, 2020; Weston et al., 2019) have been reported to have strong and selective effects at DREADDs, without pronounced off-target actions. We hope that the field will soon converge upon the “best” DREADD agonist for most behavioral experiments.

Though preliminary, these data are also valuable as proof of concept for testing DREADD agonists against one another in the same animals, in a direct and empirical manner. We hope this report will inspire others to similarly test other promising DREADD agonist drugs for their selective efficacy in head-to-head comparisons in other species and strains of model organisms and in other task conditions.

## Acknowledgements

We thank NIDA Drug Supply Program for CNO and NIMH Drug Supply Program for JHU37160.

## Data Availability Statement

Data will be made available upon request.

## Conflicts of Interest

The authors declare no conflicts of interest.

## Notes

### Competing Interest Statement

The authors have declared no competing interest.

## References

1. Alexander, G.M., Rogan, S.C., Abbas, A.I., Armbruster, B.N., Pei, Y., Allen, J.A., Nonneman, R.J., Hartmann, J., Moy, S.S., Nicolelis, M.A., McNamara, J.O., Roth, B.L., 2009. Remote control of neuronal activity in transgenic mice expressing evolved G protein-coupled receptors. Neuron 63, 27–39. https://doi.org/10.1016/j.neuron.2009.06.014

2. Armbruster, B.N., Li, X., Pausch, M.H., Herlitze, S., Roth, B.L., 2007. Evolving the lock to fit the key to create a family of G protein-coupled receptors potently activated by an inert ligand. Proc. Natl. Acad. Sci. 104, 5163–5168. https://doi.org/10.1073/pnas.0700293104

3. Atasoy, D., Sternson, S.M., 2018. Chemogenetic Tools for Causal Cellular and Neuronal Biology. Physiol. Rev. 98, 391–418. https://doi.org/10.1152/physrev.00009.2017

4. Bonaventura, J., Eldridge, M.A.G., Hu, F., Gomez, J.L., Sanchez-Soto, M., Abramyan, A.M., Lam, S., Boehm, M.A., Ruiz, C., Farrell, M.R., Moreno, A., Galal Faress, I.M., Andersen, N., Lin, J.Y., Moaddel, R., Morris, P.J., Shi, L., Sibley, D.R., Mahler, S.V., Nabavi, S., Pomper, M.G., Bonci, A., Horti, A.G., Richmond, B.J., Michaelides, M., 2019. High-potency ligands for DREADD imaging and activation in rodents and monkeys. Nat. Commun. 10, 4627. https://doi.org/10.1038/s41467-019-12236-z

5. Brodnik, Z.D., Xu, W., Batra, A., Lewandowski, S.I., Ruiz, C.M., Mortensen, O.V., Kortagere, S., Mahler, S.V., España, R.A., 2020. Chemogenetic Manipulation of Dopamine Neurons Dictates Cocaine Potency at Distal Dopamine Transporters. J. Neurosci. 40, 8767–8779. https://doi.org/10.1523/JNEUROSCI.0894-20.2020

6. Buchta, W.C., Mahler, S.V., Harlan, B., Aston-Jones, G.S., Riegel, A.C., 2017. Dopamine terminals from the ventral tegmental area gate intrinsic inhibition in the prefrontal cortex. Physiol. Rep. 5, e13198. https://doi.org/10.14814/phy2.13198

7. Burnett, C.J., Krashes, M.J., 2016. Resolving Behavioral Output via Chemogenetic Designer Receptors Exclusively Activated by Designer Drugs. J. Neurosci. 36, 9268–9282. https://doi.org/10.1523/JNEUROSCI.1333-16.2016

8. Campbell, E.J., Marchant, N.J., 2018. The use of chemogenetics in behavioural neuroscience: receptor variants, targeting approaches and caveats. Br. J. Pharmacol. 175, 994–1003. https://doi.org/10.1111/bph.14146

9. Chen, X., Choo, H., Huang, X.-P., Yang, X., Stone, O., Roth, B.L., Jin, J., 2015. The First Structure–Activity Relationship Studies for Designer Receptors Exclusively Activated by Designer Drugs. ACS Chem. Neurosci. 6, 476–484. https://doi.org/10.1021/cn500325v

10. Desloovere, J., Boon, P., Larsen, L.E., Goossens, M.-G., Delbeke, J., Carrette, E., Wadman, W., Vonck, K., Raedt, R., 2021. Chemogenetic Seizure Control with Clozapine and the Novel Ligand JHU37160 Outperforms the Effects of Levetiracetam in the Intrahippocampal Kainic Acid Mouse Model. Neurotherapeutics. https://doi.org/10.1007/s13311-021-01160-0

11. Farrell, M.R., Esteban, J.S.D., Faget, L., Floresco, S.B., Hnasko, T.S., Mahler, S.V., 2021. Ventral Pallidum GABA Neurons Mediate Motivation Underlying Risky Choice. J. Neurosci. 41, 4500–4513. https://doi.org/10.1523/JNEUROSCI.2039-20.2021

12. Farrell, M.R., Ruiz, C.M., Castillo, E., Faget, L., Khanbijian, C., Liu, S., Schoch, H., Rojas, G., Huerta, M.Y., Hnasko, T.S., Mahler, S.V., 2019. Ventral pallidum is essential for cocaine relapse after voluntary abstinence in rats. Neuropsychopharmacology 44, 2174–2185. https://doi.org/10.1038/s41386-019-0507-4

13. Ferguson, S.M., Phillips, P.E.M., Roth, B.L., Wess, J., Neumaier, J.F., 2013. Direct-Pathway Striatal Neurons Regulate the Retention of Decision-Making Strategies. J. Neurosci. 33, 11668–11676. https://doi.org/10.1523/JNEUROSCI.4783-12.2013

14. Ferrari, L.L., Ogbeide-Latario, O.E., Gompf, H.S., Anaclet, C., 2022. Validation of DREADD Agonists and Administration Route in a Murine Model of Sleep Enhancement. J. Neurosci. Methods 109679. https://doi.org/10.1016/j.jneumeth.2022.109679

15. Flerlage, W.J., Langlois, L.D., Rusnak, M., Simmons, S.C., Gouty, S., Armstrong, R.C., Cox, B.M., Symes, A.J., Tsuda, M.C., Nugent, F.S., 2022. Involvement of Lateral Habenula Dysfunction in Repetitive Mild Traumatic Brain Injury–Induced Motivational Deficits. J. Neurotrauma neu.2022.0224. https://doi.org/10.1089/neu.2022.0224

16. Fleury Curado, T., Pho, H., Freire, C., Amorim, M.R., Bonaventura, J., Kim, L.J., Lee, R., Cabassa, M.E., Streeter, S.R., Branco, L.G., Sennes, L.U., Fishbein, K., Spencer, R.G., Schwartz, A.R., Brennick, M.J., Michaelides, M., Fuller, D.D., Polotsky, V.Y., 2021. Designer Receptors Exclusively Activated by Designer Drugs Approach to Treatment of Sleep-disordered Breathing. Am. J. Respir. Crit. Care Med. 203, 102–110. https://doi.org/10.1164/rccm.202002-0321OC

17. Fortress, A.M., Hamlett, E.D., Vazey, E.M., Aston-Jones, G., Cass, W.A., Boger, H.A., Granholm, A.-C.E., 2015. Designer Receptors Enhance Memory in a Mouse Model of Down Syndrome. J. Neurosci. 35, 1343–1353. https://doi.org/10.1523/JNEUROSCI.2658-14.2015

18. Giannotti, G., Gong, S., Fayette, N., Heinsbroek, J.A., Orfila, J.E., Herson, P.S., Ford, C.P., Peters, J., 2021. Extinction blunts paraventricular thalamic contributions to heroin relapse. Cell Rep. 36, 109605. https://doi.org/10.1016/j.celrep.2021.109605

19. Gomez, J.L., Bonaventura, J., Lesniak, W., Mathews, W.B., Sysa-Shah, P., Rodriguez, L.A., Ellis, R.J., Richie, C.T., Harvey, B.K., Dannals, R.F., Pomper, M.G., Bonci, A., Michaelides, M., 2017. Chemogenetics revealed: DREADD occupancy and activation via converted clozapine. Science 357, 503–507. https://doi.org/10.1126/science.aan2475

20. Goossens, M.-G., Boon, P., Wadman, W., Van den Haute, C., Baekelandt, V., Verstraete, A.G., Vonck, K., Larsen, L.E., Sprengers, M., Carrette, E., Desloovere, J., Meurs, A., Delbeke, J., Vanhove, C., Raedt, R., 2021. Long-term chemogenetic suppression of seizures in a multifocal rat model of temporal lobe epilepsy. Epilepsia 62, 659–670. https://doi.org/10.1111/epi.16840

21. Goutaudier, R., Coizet, V., Carcenac, C., Carnicella, S., 2020. Compound 21, a two-edged sword with both DREADD-selective and off-target outcomes in rats. PLOS ONE 15, e0238156. https://doi.org/10.1371/journal.pone.0238156

22. Heinsbroek, J.A., Giannotti, G., Mandel, M.R., Josey, M., Aston-Jones, G., James, M.H., Peters, J., 2021. A common limiter circuit for opioid choice and relapse identified in a rodent addiction model. Nat. Commun. 12, 4788. https://doi.org/10.1038/s41467-021-25080-x

23. Huang, T., Guan, F., Licinio, J., Wong, M.-L., Yang, Y., 2021. Activation of septal OXTr neurons induces anxiety-but not depressive-like behaviors. Mol. Psychiatry 26, 7270–7279. https://doi.org/10.1038/s41380-021-01283-y

24. Ilg, A.-K., Enkel, T., Bartsch, D., Bähner, F., 2018. Behavioral Effects of Acute Systemic Low-Dose Clozapine in Wild-Type Rats: Implications for the Use of DREADDs in Behavioral Neuroscience. Front. Behav. Neurosci. 12.

25. Jendryka, M., Palchaudhuri, M., Ursu, D., van der Veen, B., Liss, B., Kätzel, D., Nissen, W., Pekcec, A., 2019. Pharmacokinetic and pharmacodynamic actions of clozapine-N-oxide, clozapine, and compound 21 in DREADD-based chemogenetics in mice. Sci. Rep. 9, 4522. https://doi.org/10.1038/s41598-019-41088-2

26. Kljakic, O., Hogan-Cann, A.E., Yang, H., Dover, B., Al-Onaizi, M., Prado, M.A.M., Prado, V.F., 2022. Chemogenetic activation of VGLUT3-expressing neurons decreases movement. Eur. J. Pharmacol. 935, 175298. https://doi.org/10.1016/j.ejphar.2022.175298

27. Lawson, K.A., Flores, A.Y., Hokenson, R.E., Ruiz, C.M., Mahler, S.V., 2021. Nucleus Accumbens Chemogenetic Inhibition Suppresses Amphetamine-Induced Ultrasonic Vocalizations in Male and Female Rats. Brain Sci. 11, 1255. https://doi.org/10.3390/brainsci11101255

28. Lewis, R.G., Serra, M., Radl, D., Gori, M., Tran, C., Michalak, S.E., Vanderwal, C.D., Borrelli, E., 2020. Dopaminergic Control of Striatal Cholinergic Interneurons Underlies Cocaine-Induced Psychostimulation. Cell Rep. 31, 107527. https://doi.org/10.1016/j.celrep.2020.107527

29. Li, Y., Hollis, E., 2021. Basal Forebrain Cholinergic Neurons Selectively Drive Coordinated Motor Learning in Mice. J. Neurosci. 41, 10148–10160. https://doi.org/10.1523/JNEUROSCI.1152-21.2021

30. MacLaren, D.A.A., Browne, R.W., Shaw, J.K., Krishnan Radhakrishnan, S., Khare, P., España, R.A., Clark, S.D., 2016. Clozapine N-Oxide Administration Produces Behavioral Effects in Long-Evans Rats: Implications for Designing DREADD Experiments. eNeuro 3, ENEURO.0219-16.2016. https://doi.org/10.1523/ENEURO.0219-16.2016

31. Mahler, S.V., Aston-Jones, G., 2018. CNO Evil? Considerations for the Use of DREADDs in Behavioral Neuroscience. Neuropsychopharmacology 43, 934–936. https://doi.org/10.1038/npp.2017.299

32. Mahler, S.V., Brodnik, Z.D., Cox, B.M., Buchta, W.C., Bentzley, B.S., Quintanilla, J., Cope, Z.A., Lin, E.C., Riedy, M.D., Scofield, M.D., Messinger, J., Ruiz, C.M., Riegel, A.C., España, R.A., Aston-Jones, G., 2019. Chemogenetic Manipulations of Ventral Tegmental Area Dopamine Neurons Reveal Multifaceted Roles in Cocaine Abuse. J. Neurosci. 39, 503–518. https://doi.org/10.1523/JNEUROSCI.0537-18.2018

33. Mahler, S.V., Vazey, E.M., Beckley, J.T., Keistler, C.R., McGlinchey, E.M., Kaufling, J., Wilson, S.P., Deisseroth, K., Woodward, J.J., Aston-Jones, G., 2014. Designer receptors show role for ventral pallidum input to ventral tegmental area in cocaine seeking. Nat. Neurosci. 17, 577–585. https://doi.org/10.1038/nn.3664

34. Manvich, D.F., Webster, K.A., Foster, S.L., Farrell, M.S., Ritchie, J.C., Porter, J.H., Weinshenker, D., 2018. The DREADD agonist clozapine N-oxide (CNO) is reverse-metabolized to clozapine and produces clozapine-like interoceptive stimulus effects in rats and mice. Sci. Rep. 8, 3840. https://doi.org/10.1038/s41598-018-22116-z

35. Martinez, M.X., Farrell, M.R., Mahler, S.V., 2023. Pathway-specific chemogenetic manipulation by applying ligand to axonally-expressed DREADDs, in: Vectorology for Optogenetics and Chemogenetics.

36. Martinez, V.K., Saldana-Morales, F., Sun, J.J., Zhu, P.J., Costa-Mattioli, M., Ray, R.S., 2019. Off-Target Effects of Clozapine-N-Oxide on the Chemosensory Reflex Are Masked by High Stress Levels. Front. Physiol. 10.

37. Mazzone, C.M., Pati, D., Michaelides, M., DiBerto, J., Fox, J.H., Tipton, G., Anderson, C., Duffy, K., McKlveen, J.M., Hardaway, J.A., Magness, S.T., Falls, W.A., Hammack, S.E., McElligott, Z.A., Hurd, Y.L., Kash, T.L., 2018. Acute engagement of Gq-mediated signaling in the bed nucleus of the stria terminalis induces anxiety-like behavior. Mol. Psychiatry 23, 143–153. https://doi.org/10.1038/mp.2016.218

38. Nagai, Y., Miyakawa, N., Takuwa, H., Hori, Y., Oyama, K., Ji, B., Takahashi, M., Huang, X.-P., Slocum, S.T., DiBerto, J.F., Xiong, Y., Urushihata, T., Hirabayashi, T., Fujimoto, A., Mimura, K., English, J.G., Liu, J., Inoue, K., Kumata, K., Seki, C., Ono, M., Shimojo, M., Zhang, M.-R., Tomita, Y., Nakahara, J., Suhara, T., Takada, M., Higuchi, M., Jin, J., Roth, B.L., Minamimoto, T., 2020. Deschloroclozapine, a potent and selective chemogenetic actuator enables rapid neuronal and behavioral modulations in mice and monkeys. Nat. Neurosci. 23, 1157–1167. https://doi.org/10.1038/s41593-020-0661-3

39. Nentwig, T.B., Obray, J.D., Vaughan, D.T., Chandler, L.J., 2022. Behavioral and slice electrophysiological assessment of DREADD ligand, deschloroclozapine (DCZ) in rats. Sci. Rep. 12, 6595. https://doi.org/10.1038/s41598-022-10668-0

40. O’Neal, T.J., Nooney, M.N., Thien, K., Ferguson, S.M., 2020. Chemogenetic modulation of accumbens direct or indirect pathways bidirectionally alters reinstatement of heroin-seeking in highbut not low-risk rats. Neuropsychopharmacology 45, 1251–1262. https://doi.org/10.1038/s41386-019-0571-9

41. Paxinos, G., Watson, C., 2006. The Rat Brain in Stereotaxic Coordinates: Hard Cover Edition. Elsevier.

42. Porter, J.H., Manvich, D.F., Webster, K.A., Foster, S.L., Farrell, M.S., Weinshenker, D., 2017. Behavioral and Pharmacokinetic Properties of the Putatively-Inert DREADD Ligand Clozapine-N-Oxide (CNO) in Rats and Mice. FASEB J. 31, lb594–lb594.

43. Raper, J., Galvan, A., 2022. Applications of chemogenetics in non-human primates. Curr. Opin. Pharmacol. 102204. https://doi.org/10.1016/j.coph.2022.102204

44. Raper, J., Morrison, R.D., Daniels, J.S., Howell, L., Bachevalier, J., Wichmann, T., Galvan, A., 2017. Metabolism and Distribution of Clozapine-N-oxide: Implications for Nonhuman Primate Chemogenetics. ACS Chem. Neurosci. 8, 1570–1576. https://doi.org/10.1021/acschemneuro.7b00079

45. Rinker, J.A., Marshall, S.A., Mazzone, C.M., Lowery-Gionta, E.G., Gulati, V., Pleil, K.E., Kash, T.L., Navarro, M., Thiele, T.E., 2017. Extended Amygdala to Ventral Tegmental Area Corticotropin-Releasing Factor Circuit Controls Binge Ethanol Intake. Biol. Psychiatry, The Extended Amygdala and Addiction 81, 930–940. https://doi.org/10.1016/j.biopsych.2016.02.029

46. Rogan, S.C., Roth, B.L., 2011. Remote Control of Neuronal Signaling. Pharmacol. Rev. 63, 291–315. https://doi.org/10.1124/pr.110.003020

47. Rorabaugh, J.M., Chalermpalanupap, T., Botz-Zapp, C.A., Fu, V.M., Lembeck, N.A., Cohen, R.M., Weinshenker, D., 2017. Chemogenetic locus coeruleus activation restores reversal learning in a rat model of Alzheimer’s disease. Brain 140, 3023–3038. https://doi.org/10.1093/brain/awx232

48. Salimi-Nezhad, N., Missault, S., Reinoso, A.N., Hassani, A., Amiri, M., Keliris, G.A., 2023. The impact of selective and non-selective medial septum stimulation on hippocampal neuronal oscillations: A study based on modeling and experiments. Neurobiol. Dis. 106052. https://doi.org/10.1016/j.nbd.2023.106052

49. Smith, K.S., Bucci, D.J., Luikart, B.W., Mahler, S.V., 2016. DREADDs: Use and Application in Behavioral Neuroscience. Behav. Neurosci. 130, 137–155. https://doi.org/10.1037/bne0000135

50. Song, J., Patel, R.V., Sharif, M., Ashokan, A., Michaelides, M., 2022. Chemogenetics as a neuromodulatory approach to treating neuropsychiatric diseases and disorders. Mol. Ther. 30, 990–1005. https://doi.org/10.1016/j.ymthe.2021.11.019

51. Stachniak, T.J., Ghosh, A., Sternson, S.M., 2014. Chemogenetic Synaptic Silencing of Neural Circuits Localizes a Hypothalamus→Midbrain Pathway for Feeding Behavior. Neuron 82, 797–808. https://doi.org/10.1016/j.neuron.2014.04.008

52. Thompson, K.J., Khajehali, E., Bradley, S.J., Navarrete, J.S., Huang, X.P., Slocum, S., Jin, J., Liu, J., Xiong, Y., Olsen, R.H.J., Diberto, J.F., Boyt, K.M., Pina, M.M., Pati, D., Molloy, C., Bundgaard, C., Sexton, P.M., Kash, T.L., Krashes, M.J., Christopoulos, A., Roth, B.L., Tobin, A.B., 2018. DREADD Agonist 21 Is an Effective Agonist for Muscarinic-Based DREADDs in Vitro and in Vivo. ACS Pharmacol. Transl. Sci. 1, 61–72. https://doi.org/10.1021/acsptsci.8b00012

53. Upright, N.A., Baxter, M.G., 2020. Effect of chemogenetic actuator drugs on prefrontal cortex-dependent working memory in nonhuman primates. Neuropsychopharmacology 45, 1793–1798. https://doi.org/10.1038/s41386-020-0660-9

54. Urban, D.J., Roth, B.L., 2015. DREADDs (Designer Receptors Exclusively Activated by Designer Drugs): Chemogenetic Tools with Therapeutic Utility. Annu. Rev. Pharmacol. Toxicol. 55, 399–417. https://doi.org/10.1146/annurev-pharmtox-010814-124803

55. Weston, M., Kaserer, T., Wu, A., Mouravlev, A., Carpenter, J.C., Snowball, A., Knauss, S., von Schimmelmann, M., During, M.J., Lignani, G., Schorge, S., Young, D., Kullmann, D.M., Lieb, A., 2019. Olanzapine: A potent agonist at the hM4D(Gi) DREADD amenable to clinical translation of chemogenetics. Sci. Adv. 5, eaaw1567. https://doi.org/10.1126/sciadv.aaw1567

56. Whissell, P.D., Tohyama, S., Martin, L.J., 2016. The Use of DREADDs to Deconstruct Behavior. Front. Genet. 7.

57. Zhang, J., Chen, D., Sweeney, P., Yang, Y., 2020. An excitatory ventromedial hypothalamus to paraventricular thalamus circuit that suppresses food intake. Nat. Commun. 11, 6326. https://doi.org/10.1038/s41467-020-20093-4

58. Zhu, H., Roth, B.L., 2015. DREADD: A Chemogenetic GPCR Signaling Platform. Int. J. Neuropsychopharmacol. 18, pyu007. https://doi.org/10.1093/ijnp/pyu007

